# The perils of interaction prediction

**DOI:** 10.1101/435065

**Authors:** Weiguang Mao, Dennis Kostka, Maria Chikina

**Affiliations:** Department of Computational and Systems Biology, University of Pittsburgh, Pittsburgh, PA, USA; Joint Carnegie Mellon University-University of Pittsburgh Ph.D. Program in Computational Biology, Pittsburgh, PA, USA; Department of Developmental Biology, School of Medicine, University of Pittsburgh, Pittsburgh, PA, USA

## Abstract

The availability of genome-wide maps of enhancer-promoter interactions (EPIs) has made it possible to use machine learning approaches to extract and interpret features that determine these interactions in different biological contexts. Multiple methods have claimed to accomplish the task of predicting enhancer-promoter interactions based on corresponding genomic features, but this problem is actually still far from being solved. In our analysis, we show that individual enhancer and promoter regions have widely different marginal interaction probabilities, e.g. propensities, which can lead to overfitting and memorization when random cross-validation is employed. Further even when a proper cross-validation scheme is adopted, a simple propensity-based model can still achieve a competitive performance without capturing any information about the EPI mechanism.

## Introduction

Eukaryotic gene regulation is a highly complex process that involves the interaction of distal cis-regulatory elements, like enhancers, with the promoters of their target genes involving multi-protein complexes. The current state of the art to assay such enhancer-promoter interactions (EPIs) is to perform high-throughput chromatin conformation capture (Hi-C), which is an expensive technology requiring large numbers of cells. Therefore, there is a great interest in whether computational approaches can predict EPIs from other assays or genomic annotations, including from DNA sequences alone. In recent years there has been a proliferation of methods that claim to accomplish this task [1, 2, 3]. Since these methods are not able to make truly novel predictions (*i.e.*, performance in an unseen biological context is poor), the focus has been on interpreting the models themselves [1]. The hope is that analyzing which models work best and which features are predictive may give us some insight into the mechanistic aspects of EPIs.

Unfortunately, in our analysis we show that for the gold standard of EPIs used in several previous studies, it is possible to achieve a very good performance without learning *anything* about what drives specific interactions. Instead, the models learn to predict the interaction “propensity” of individual sequence elements, which is defined as the marginal probability that a particular sequence element interacts with *any other* in a given gold standard. The final interaction probability can be computed as a function of individual propensities giving very good performance while not capturing any information about the mechanisms of individual interactions.

## Results

We analyze the datasets compiled originally by the authors of TargetFinder which claimed to predict EPIs from various chromatin features [1]. The same standard was later used by multiple studies to claim that it is possible to make predictions with high accuracy using sequence features alone [2, 3]. This standard is heavily imbalanced in the sense that there are many promoters and enhancer for which have only positive or only negative interactions. We refer to the marginal probability that a given promoter or enhancer sequence participates in a positive interaction as its interaction propensity. For the TargetFinder gold standard these range from 0 to 1. We then evaluate how well we can recapitulate the gold standard given only the propensities and find that if the propensities are known exactly we can achieve > 99% accuracy (AUC=1) on all cell-lines with simple multiplication.

Of course, the analysis just described doesn’t account for the effect of cross validation, and the machine learning models from [1, 2, 3] were trained using 10-fold random holdout cross validation. Cross validation in our setting has a two fold effect. Instead of having the exact dataset-wide propensities we now have only estimates, and, more importantly, since some promoters and enhancers are never seen, their propensities are unknown (and just set to the average). In this 10-fold cross-validation framework we combine the propensities using two different methods: a simple multiplication and Gaussian Process classification. While the product assumes that propensities are independent, the Gaussian process can fully account for all dependencies and empirically gives better performance. But we emphasize that it is still a trivial approach designed only to estimate individual propensities for interaction.

Nevertheless this naíve method still achieves excellent performance with all AUCs above 0.93. In fact, our results are competitive with the models that use the full set of features, performing nearly as well to sequence-only PEP and only slightly worse than chromatin-feature based TargetFinder. Thus, our simple propensity model is capable of achieving performance akin to that of complex models while using *no features*. The only information our model uses are the individual region propensity estimates and the ability to match regions across the train and test split.

One issue with this analysis is that neither the original TargetFinder and PEP methods nor this propensity based analysis was done using strict cross-validation, where a set of promoters and enhancers is entirely held out of the training. Instead the methods use random cross-validation, where only pairs are held out. In this setting it is indeed possible for a machine learning approach to memorize the propensities of individual regions and our analysis here simply gives a lower bound on the performance that should be achievable from memorization alone.

It is more natural to ask how well these models perform in a strict cross-validation setting where an entire chromosome is held-out and propensity memorization is not possible. We consequently reran both TargetFinder and PEP with chromosome holdout. The performance of both of these methods decreases dramatically from what was reported (see Figure 2 circles for TargetFinder with default parameters).

**Figure 1:**
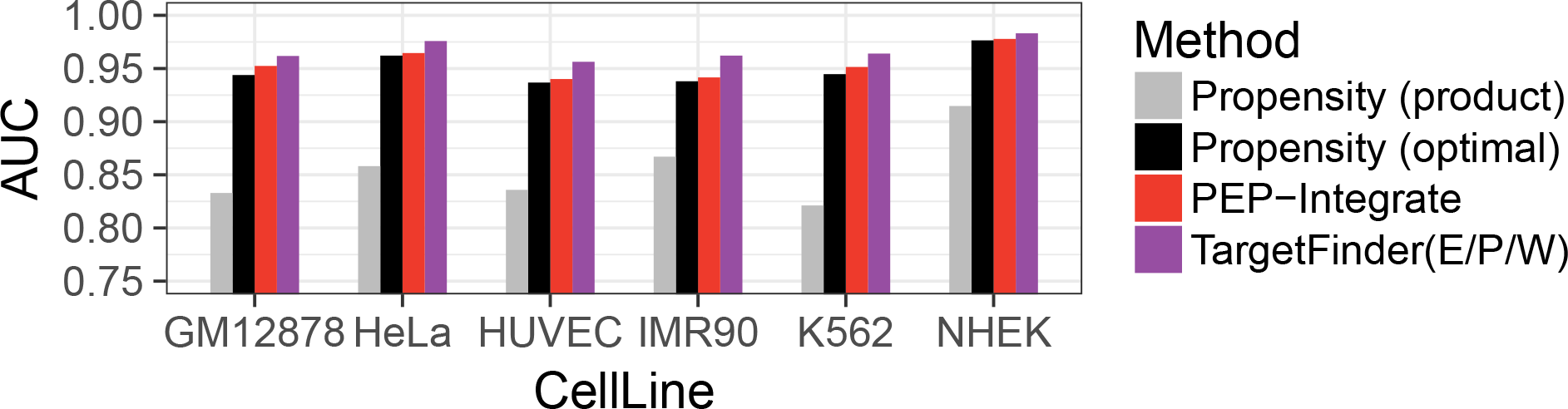
Comparing propensity based model to other machine learning methods using random holdout. Performance of PEP and TargetFinder is as reported in [2]. Propensity based model simply computes the propensity for each enhancer and promoter based on the training set. If the particular enhancer or promoter has not been observed it is assigned a the average propensity. The final EPI predictions are then generated either by simple propensity multiplication (product) or by a Gaussian Process classification with the two propensities as input (optimal).

**Figure 2:**
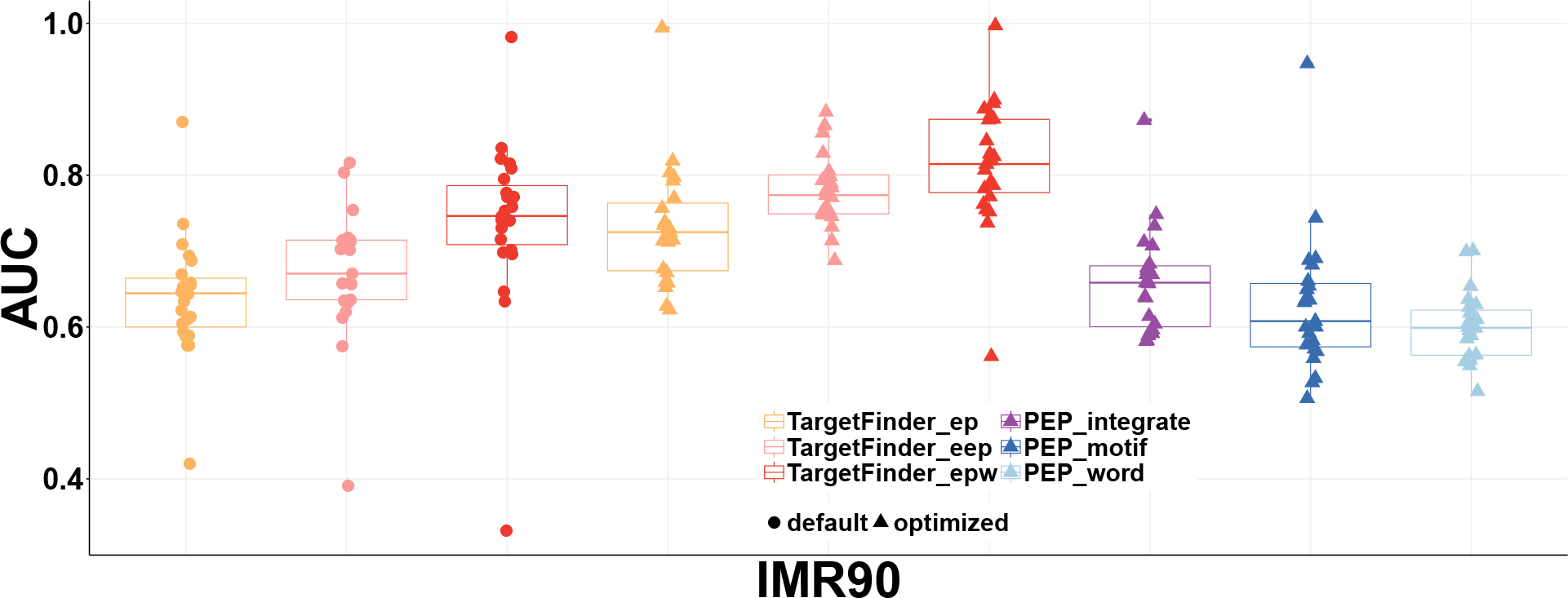
AUC metrics under different parameter settings using chromosomes held-out cross-validation based on IMR90 cell line. Models were trained with chromosome hold-out and each point stands for the test performance when the corresponding single chromosome (chr1-chrX) is held out as the test set. TargetFinder was evaluated with original reported parameters (circles) and parameters re-optimized for chromosome holdout performance (triangles). For PEP, only re-optimized performance is shown since the original parameters lead to a near random performance.

**Figure 3:**
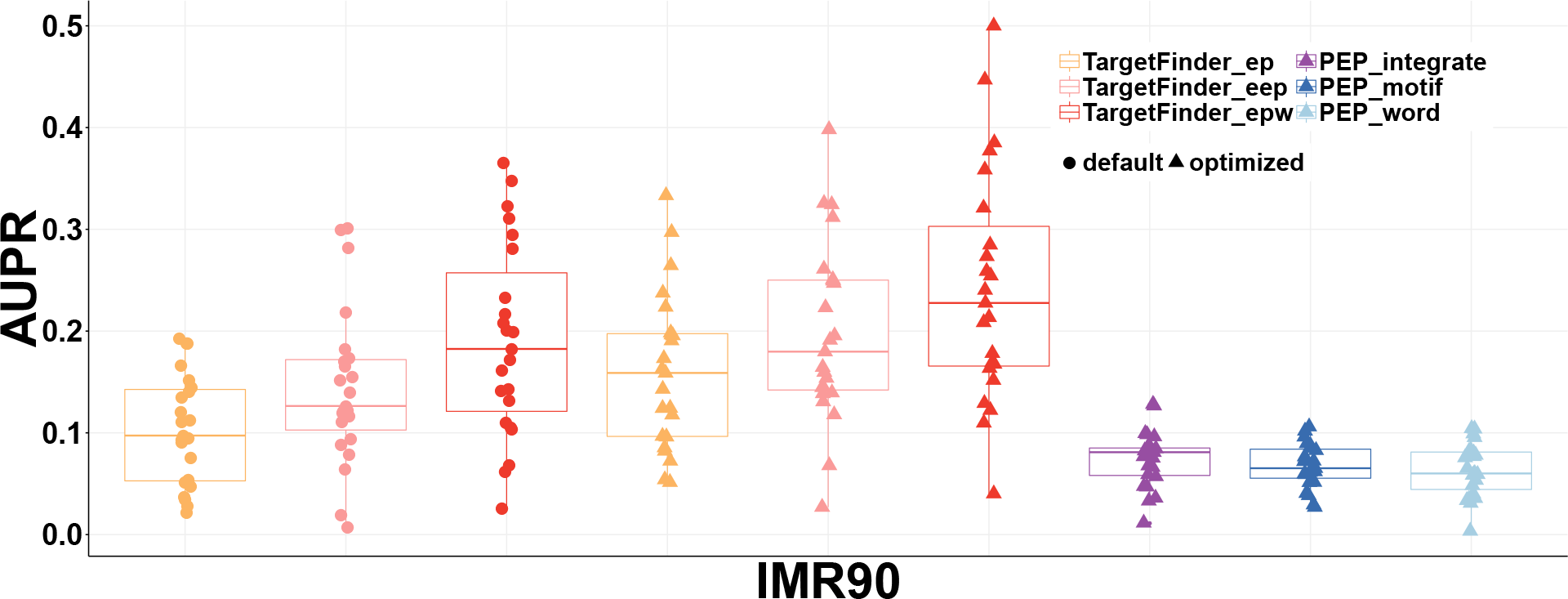
Same predictions as in Figure 2 evaluated for Area Under Precision Recall (AUPR)

Both TargetFinder and PEP rely on boosted decision trees and the published hyper-parameters were optimized using random cross validation. We thus re-optimize the hyper-parameters to maximize performance on chromosome holdout. We find that the re-optimized parameters correspond to considerably lower model capacity (see Table 1) but do improve the chromosome hold-out performance (see Figure 2 triangles) which is what would be expected if random holdout results in memorization and overfitting. We find that re-optimized PEP achieves performance that is above random and the re-optimized TargetFinder performance is reasonably good (median AUC is 0.82) though still considerably lower than originally reported.

**Table 1:** The optimized parameters for the Gradient Boosting Trees using each chromosome as the held-out test set when running each single TargetFinder model. The default parameters for the Gradient Boosting Trees in TargetFinder models are n estimators = 4000 and max depth = 5. The first number in each entry represents the number of estimators and the second number denotes the depth. In aggregate the re-optimized parameters correspond to models with lower capacity.

|  | TargetFinder_EP_optimized | TargetFinder_EEP_optimized | TargetFinder_EPW_optimized |
| --- | --- | --- | --- |
| chr1 | 20,2 | 50,2 | 200,2 |
| chr2 | 100,3 | 200,3 | 200,5 |
| chr3 | 100,5 | 500,3 | 500,2 |
| chr4 | 100,5 | 1000,2 | 500,6 |
| chr5 | 50,5 | 100,2 | 100,2 |
| chr6 | 100,2 | 50,4 | 100,3 |
| chr7 | 50,3 | 200,2 | 100,5 |
| chr8 | 1000,2 | 20,6 | 500,2 |
| chr9 | 50,2 | 50,5 | 200,2 |
| chr10 | 50,4 | 20,5 | 50,6 |
| chr11 | 20,5 | 200,4 | 200,3 |
| chr12 | 200,2 | 200,3 | 100,5 |
| chr13 | 50,6 | 100,4 | 200,3 |
| chr14 | 4000,4 | 50,5 | 500,5 |
| chr15 | 50,4 | 50,3 | 50,4 |
| chr16 | 200,3 | 100,6 | 50,6 |
| chr17 | 20,6 | 100,2 | 100,6 |
| chr18 | 100,5 | 50,4 | 200,2 |
| chr19 | 20,2 | 20,2 | 20,5 |
| chr20 | 2000,4 | 1000,4 | 500,5 |
| chr21 | 500,4 | 50,3 | 4000,2 |
| chr22 | 500,2 | 100,5 | 500,2 |
| chrX | 100,6 | 4000,4 | 200,3 |

At first glance, it appears that with proper cross validation both chromatin states and even sequence-only features contain some signal that may be used to predict EPIs. In particular, TargetFinder performance is quite good given that the EPI prediction has some minimal irreducible error due to measurement noise in both features and labels. However, given our earlier observation that propensities alone are highly predictive, the baseline level for this classification task is not in facts AUC=0.5 but the AUC that can be achieved with a propensity only model. We thus apply a propensity model in the stringent chromosome hold-out setting. In this setting the propensities are not memorized but are predicted from the TargetFinder features using standard machine learning methods.

The complete propensity based model is built in two layers (see Figure 4). In the first layer we use Gradient Boosting trees to predict propensities of individual elements (promoters, enhancers, and window regions). The single numerical result from the first layer is passed to the second layer which uses Gaussian process classifier to integrate two (E, P) or three (E, P and W) propensity values into a single interaction probability. In contrast to the TargetFinder model, this propensity based model doesn’t allow for the actual features from the E, P and W regions to interact. What are visible to the second layer are just the propensity values.

**Figure 4:**
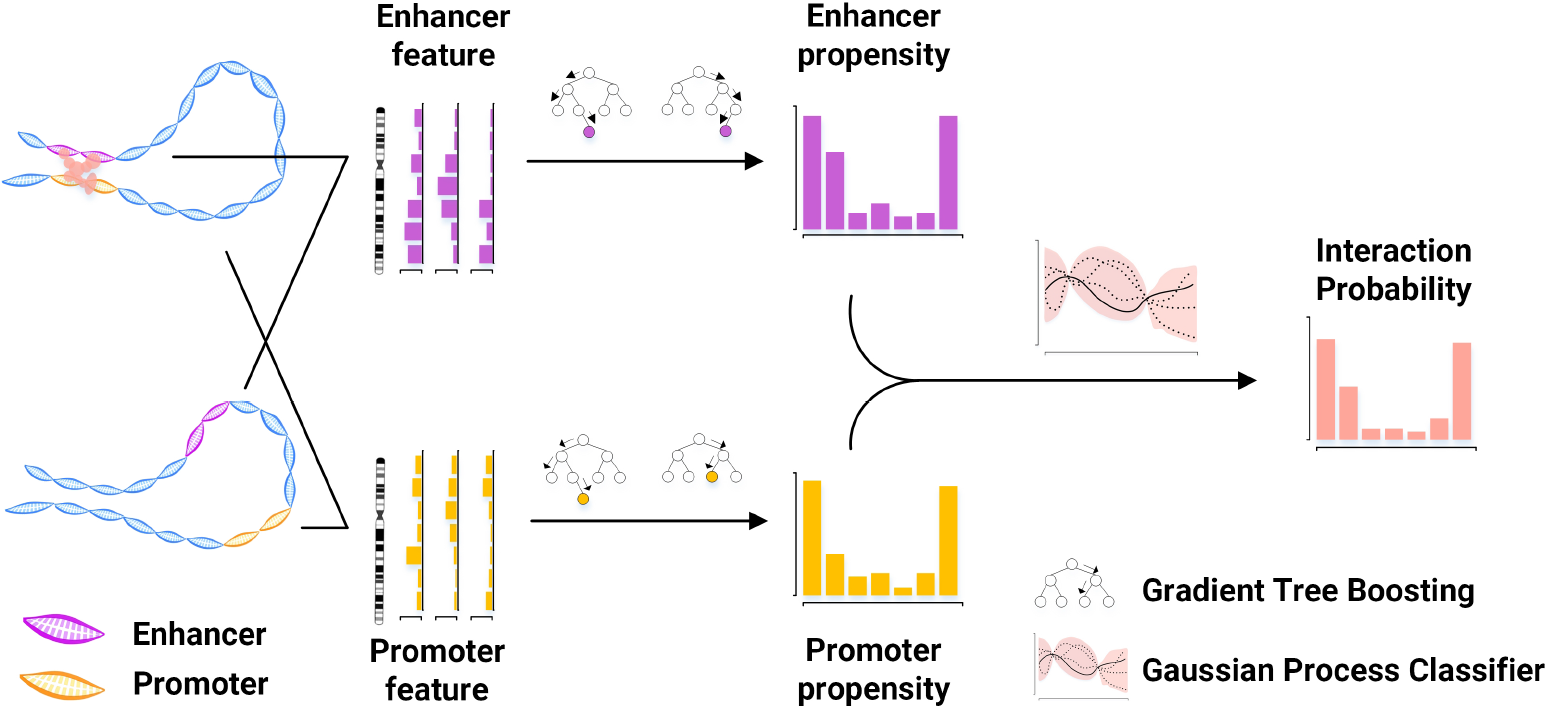
Schematic diagram of propensity model. In the first layer the promoter, enhancer, and window (not shown) features are used to build a propensity model using boosted decision trees. In the second layer the propensity values are combined into a single interaction prediction using a Gaussian Process classifier

We find that using chromosome hold-out cross-validation, the propensity-only model achieves performance that is very close to that of TargetFinder (Figure 5). Since in the propensity model the individual features from different regions do not interact, this gives us a baseline performance against which other models can be evaluated and we find that relative to this baseline the performance gains achieved by TargetFinder are minimal. In order to characterize the contribution of individual promoter, enhancer, or window regions to final performance, we also evaluate individual propensities. In this setting we simply assign the interaction probability for any EPI pair as the propensity of the individual promoter, enhance, or window element. We find that in this setting window features are most predictive though on average the complete “EPW” model is slightly better. This finding is consistent with the importance of window features highlighted in the original TargetFinder paper though our analysis demonstrates that the effect is almost entirely mediated by the window propensity.

**Figure 5:**
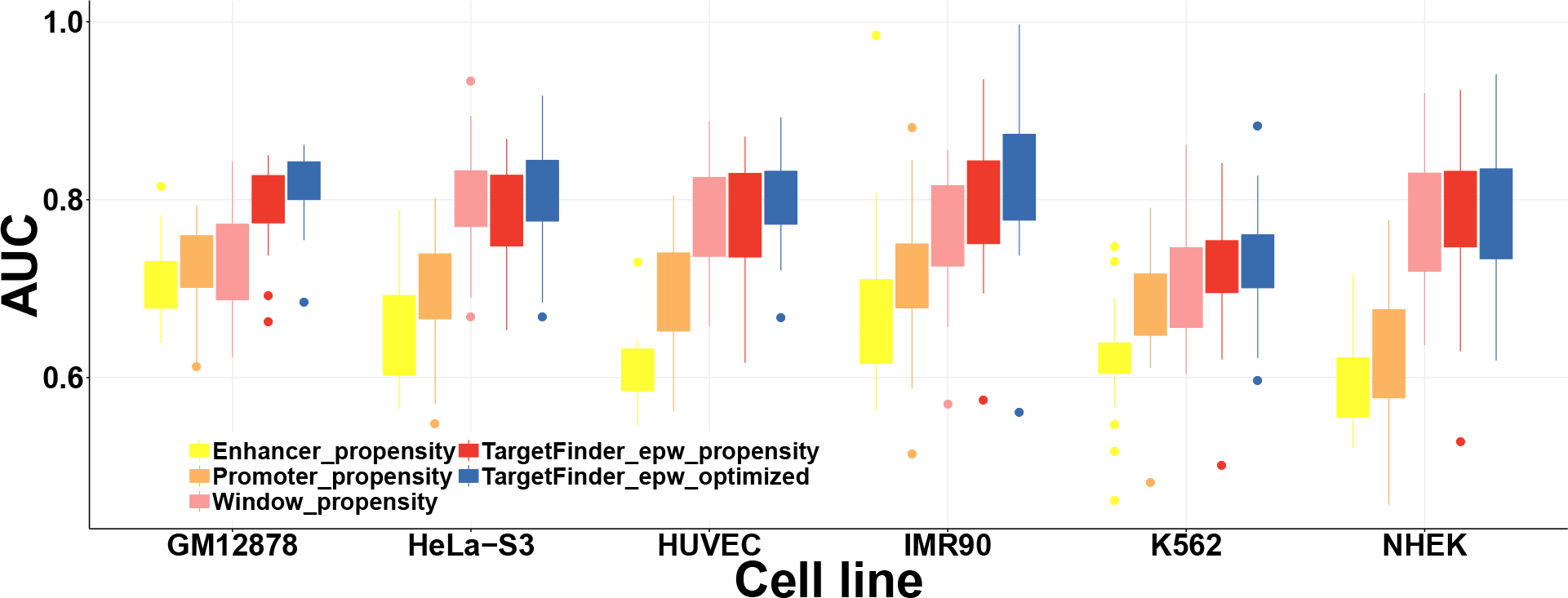
AUC metrics of different EPI prediction strategies using chromosomes held-out cross-validation across all cell lines. Enhancer propensity, promoter propensity or window propensity means that the model only passes enhancer, promoter or window propensity to evaluate the test performance.

**Figure 6:**
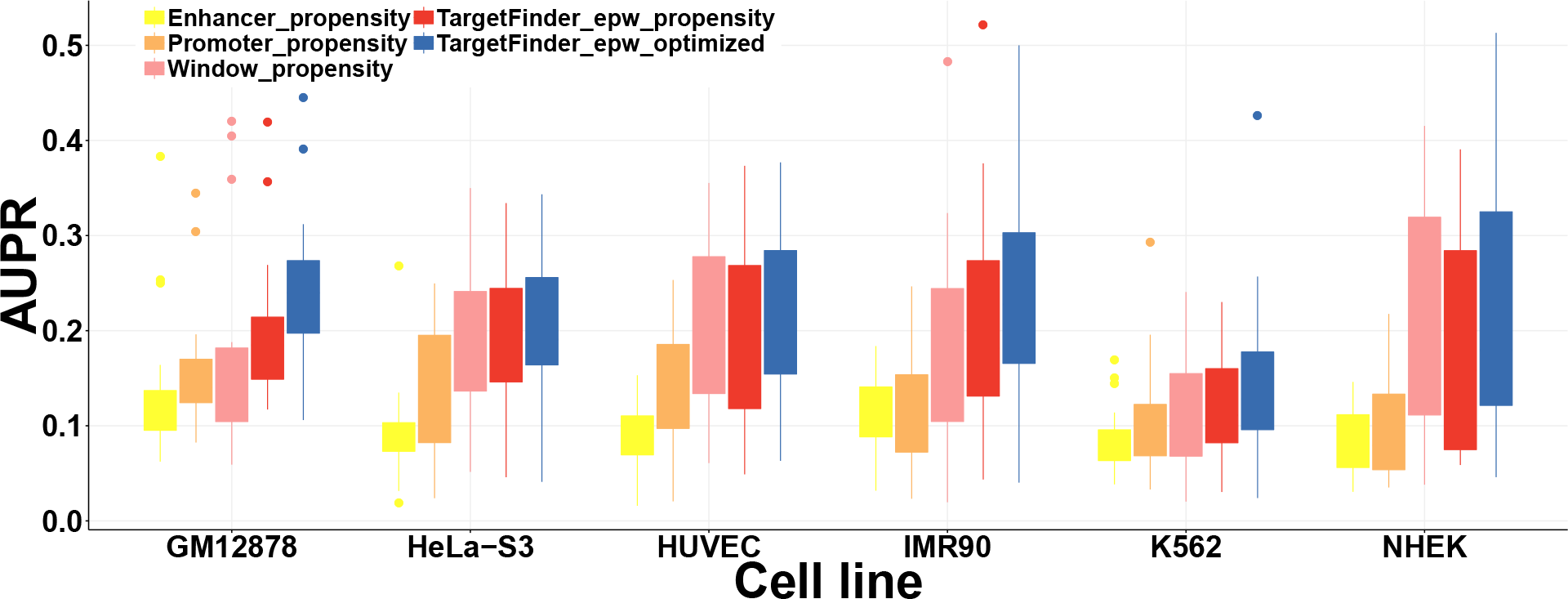
AUPR metrics of different EPI prediction strategies using chromosomes held-out cross-validation across all cell lines. Enhancer propensity, promoter_propensity or window_propensity means that the model only passes enhancer, promoter or window propensity to evaluate the test performance.

## Conclusion

In summary we demonstrate two things: firstly, random cross-validation employed for EPI prediction on the TargetFinder dataset obscures an important problem. Since the individual enhancer and promoter regions have widely different marginal interaction probabilities (referred to as interaction propensities), it is possible for machine learning methods to simply memorize these values and this alone is enough to produce a good classification performance. Our second point is that the same issue that is able to produce dramatic overfitting on random cross-validation is still at play even when proper cross-validation is performed. This is a general feature of any interaction prediction task where the marginal interaction probabilities are different from baseline. We emphasize that for such imbalanced datasets it is critical to establish the baseline predictive performance of a propensity based model. Subsequently, models that are computed on the entire feature set can be compared against that baseline. Unfortunately, for the TargetFinder dataset the propensity based model gives a very good performance leaving little room for improvement.

## Methods

### Data

We downloaded the TargetFinder data and script from the referened github repository (https://github.com/shwhalen/targetfinder) and followed the guide to rerun the computations and set up runs to optimize parameters in the chromosome held-out setting by grid search. The number of estimators is selected from [20, 50, 100, 200, 500, 1000, 2000, 4000] and the depth of the trees ranges from 2 to 6. We downloaded the PEP data and script from the github repository (https://github.com/ma-compbio/PEP) and also optimized the chromosome held-out performance by grid search. The number of estimators is selected from [20, 50, 100, 200, 500, 1000, 1500, 2000] and the depth of the trees is picked up from [2, 3, 4, 5, 10, 15].

### Propensity model

Using the re-optimized parameters we observed at that of the available TargetFinder settings (EPW, EP, and EEP) EPW achieves the best and for our propensity model we use all three features sets, which are enhancer features, promoter features and window features. In the first layer, we adopted Gradient Boosting trees to generate propensity values for each category of features. In order to constrain the complexity of the model, three Gradient Boosting classifiers share the same parameters and we optimized the number of estimators within the set [20, 50, 100, 200, 500, 1000] and the depth of tress within the set [2, 3, 4, 5, 6]. We utilized Gaussian Process classifier with the default RBF kernel and we didn’t optimize on the hyper-parameters of the RBF kernel. We trained the propensity model in a heuristic manner. We trained the first layer with enhancer feature *x*_*E,train*_, promoter feature *x*_*P,train*_, or window feature *x*_*W,train*_ and the interaction labels *y*_*train*_. Then we fed the predicted propensities *p*_*E,train*_, *p*_*P,train*_ and *p*_*W,train*_ on this train set and *y*_*train*_ into the Gaussian Process Classifier (GP). Because of the scale of the data, we only subsampled 1,000 pairs to train the GP.

